# No influence of antibiotic on locomotion in *Drosophila nigrosparsa* after recovery, but influence on microbiome, possibly mediating wing-morphology change

**DOI:** 10.1101/2021.03.09.434651

**Authors:** Simon O. Weiland, Matsapume Detcharoen, Birgit C. Schlick-Steiner, Florian M. Steiner

## Abstract

Antibiotics, such as tetracycline, has been frequently used to cure endosymbiont *Wolbachia* in arthropods. After the symbionts had been removed, the hosts must be waited for some generations to recover from side effects of the antibiotics. Knowledge of potential long-term effects of the antibiotic is important. Here, we treated *Drosophila nigrosparsa* with and without antibiotic tetracycline for three generations and two generations recovering time to investigate effects of the tetracycline on the flies concerning locomotion of larvae and adults, wing morphology, and gut microbiome of adults. In addition, gut-microbiome restoration was tested as a solution to reducing potential side effects of tetracycline on the flies’ microbiome more quickly. We found significant differences in larval and adult locomotion within groups but no significant differences among the control, antibiotic-treated, and gut-restoration groups. We found a slight differentiation of wing morphology into the three groups and significant differences in bacterial abundance among groups. The influence of tetracycline on the gut microbiome may have contributed to wing-morphology differences among groups, which would be an indirect effect of the antibiotic. Together with the absence of an effect on locomotion, this suggests that checking for both direct and indirect effects of tetracycline after a particular recovery time before using tetracycline curing is important. The microbiome of the gut-restoration group was not like that of the control group. Therefore, gut restoration cannot be used to remove effects of tetracycline in *D. nigrosparsa*, at least in the setup used here.

## INTRODUCTION

The intracellular Alphaproteobacteria *Wolbachia* belong to the Rickettsiales group (Lo *et al.*, 2007). They are found in 40-60% of arthropods (Zug & Hammerstein, 2012; Sazama *et al.*, 2017), but there is still a huge knowledge gap about *Wolbachia* diversity, as less than 1% of the expected 100,000 strains in arthropods is known (Detcharoen *et al.*, 2019). They are usually maternally transmitted, but horizontal transfer also occurs (Schuler *et al.*, 2013). The four major phenotypic effects of *Wolbachia* on their hosts are cytoplasmic incompatibility, male killing, parthenogenesis, and feminization, of which cytoplasmic incompatibility is the most studied (Werren *et al.*, 2008). *Wolbachia* can also induce other changes; for example, infected *Drosophila melanogaster* can have a higher mating rate (De Crespigny *et al.*, 2006), higher fecundity (Fry *et al.*, 2004; Serga *et al.*, 2014), more resistance to viruses (Teixeira *et al.*, 2008; Chrostek *et al.*, 2013), and morphological changes such as larger wings (Kriesner *et al.*, 2016). To pinpoint *Wolbachia* as causal to a suspected change in a host, establishing a comparison group by curing infected hosts from *Wolbachia* is necessary. One of the most frequently used solutions for curing is a tetracycline treatment (Schneider *et al.*, 2013).

Tetracycline is a broad-spectrum antibiotic and inhibits the prokaryotic 30S ribosomes. It is used against various bacterial infections (Ballard & Melvin, 2007), but it also acts on host enzymes and mitochondrial proteins by inhibiting the metabolism, synthesis, and repair of nucleic acids (Brodersen *et al.*, 2000). In *Drosophila*, tetracycline continues affecting mitochondrial DNA density, mitochondrial metabolism (Ballard & Melvin, 2007), and the host’s fitness after the treatment has ceased (Miller *et al.*, 2010). Therefore, after the antibiotic treatment, a recovery time before starting further experiments is important. Two generations have been reported as sufficient in *Drosophila* to reduce side effects from tetracycline (Fry *et al.*, 2004; Harcombe & Hoffmann, 2004).

The composition of gut microbiota also changes with the antibiotic treatment (Raymann *et al.*, 2017; Jung *et al.*, 2018). Changes in the abundance of Proteobacteria and Firmicutes in the microbiome after tetracycline treatment were described (Chao *et al.*, 2020). In addition, several studies have identified importance of various bacterial taxa on their hosts, such as development (Buchon *et al.*, 2009; Storelli *et al.*, 2011), physiological processes, lifespan (Sommer & Bäckhed, 2013; Gilbert *et al.*, 2018), disease resistance (Sansone *et al.*, 2015), and behavior (Selkrig *et al.*, 2018).

The aim of this study is to examine the effect of tetracycline on *Drosophila nigrosparsa* two generations after using tetracycline. Better knowing the long-term effects of tetracycline on *D. nigrosparsa* is needed for better interpreting published (Detcharoen *et al.*, 2020) and future results, given that this species is in the focus of climate-change research (Kinzner *et al.*, 2019). Thus, we ask if changes in the flies result from previous *Wolbachia* infection or previous tetracycline treatment. For this purpose, non-infected *D. nigrosparsa* are treated with tetracycline and compared with a control group to investigate the sole effect of the antibiotic on the flies. The experiments examine the locomotion of larvae and adults as well as the wing morphology, which were included in a previous study examining the effects of *Wolbachia* on *D. nigrosparsa* (Detcharoen *et al.*, 2020). The gut microbiome is characterized in this study, and in addition, gut-microbiome restoration is tested as a solution to reducing potential side effects of tetracycline on the microbiome more quickly than without using it.

## MATERIAL & METHODS

### Study system *Drosophila nigrosparsa*

The distribution area of *D. nigrosparsa* is Central and Western Europe. In Central Europe (Bächli *et al.*, 1985, 2004), the fly lives at about 2000 m above sea level (Bächli *et al.*, 1985, 2004). In various physiological limits and life history traits, the fly is well adapted to its extreme environment (Kinzner *et al.*, 2016, 2018; Tratter Kinzner *et al.*, 2019). Under artificial conditions at 19 °C, the development time (embryo to adult) is around 60 days (Kinzner *et al.*, 2016). No natural infection of *Wolbachia* in *D. nigrosparsa* is known (Detcharoen *et al.*, 2020). Detcharoen et al. (2020) transinfected *D. nigrosparsa* with the *Wolbachia* strain *w*Mel and cured it with tetracycline to study the effects of *w*Mel on the flies.

### Fly lines

*Drosophila nigrosparsa* was collected at Kaserstattalm in Stubai Valley, Tyrol, Austria (47.13°N, 11.30°E) in 2010 (Kinzner *et al.*, 2018) and used to establish the isofemale line iso12 (Arthofer *et al.*, 2015; Cicconardi *et al.*, 2017). The isofemale line used in this study was a subpopulation of iso12 and was used in previous studies (Detcharoen *et al.*, 2020). It was used to establish three control lines (not treated with antibiotic), namely, −T1, −T2, and −T3, and three antibiotic-treated lines, namely, +T1, +T2, and +T3 (Figure 1). Gut restoration lines, +T11, +T22, and +T33 were created by splitting the antibiotic-treated lines in Generation 5. All flies were kept in mating cages (50 adult females and 50 adult males) (Kinzner *et al.*, 2018) and supplied with grape juice agar, malt food, and yeast. Food was changed twice a week. Embryos or first-stage larvae were transferred to glass vials with 8 ml malt food at a density of 80 embryos or 60 larvae per vial, respectively. All flies were kept at 19 °C, 70% humidity, and a 16h:8h light:dark cycle.

**Figure 1.**
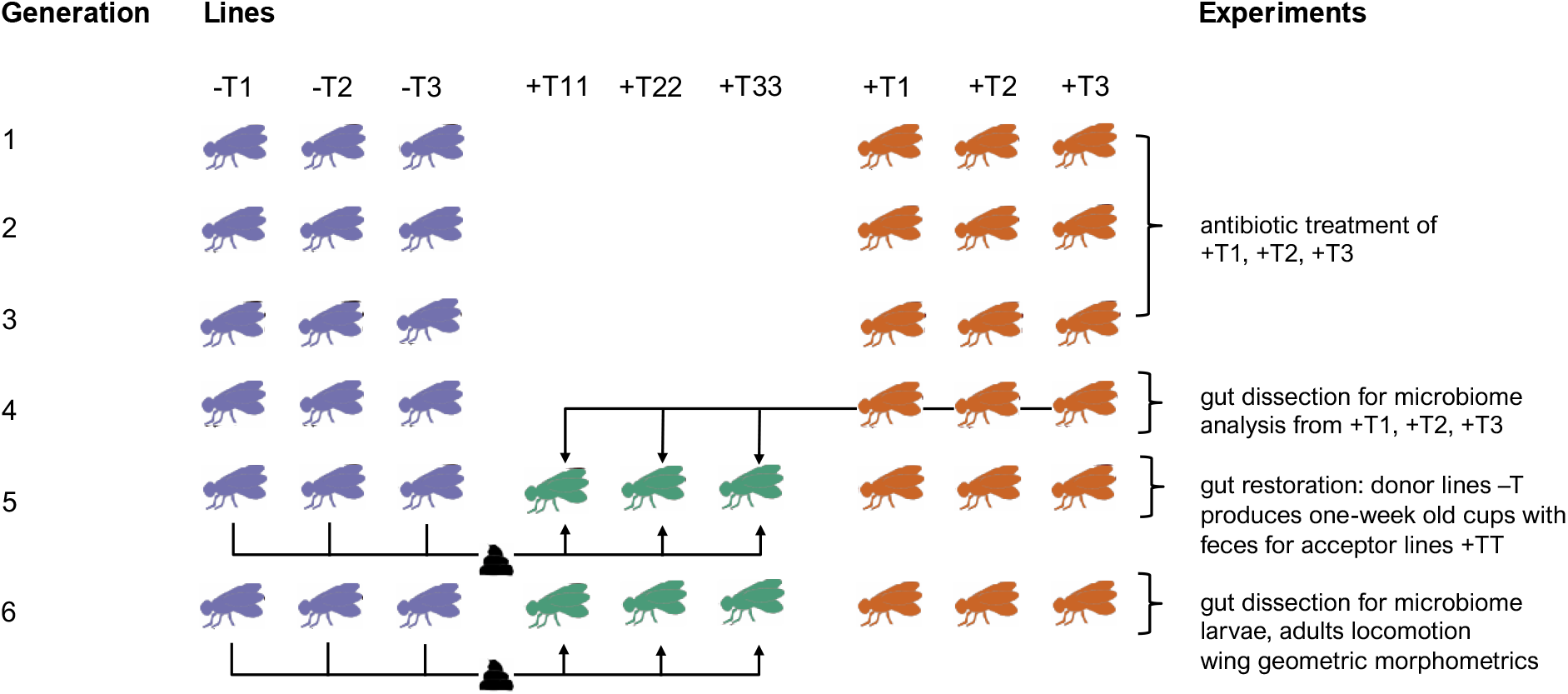
Chronological overview of the study using *Drosophila nigrosparsa*. Each fly line was kept in mating cages at a census size of 50 males and 50 females in every generation. Control lines −T1, −T2, and −T3 are fly lines not treated with tetracycline. Antibiotic-treated lines +T1, +T2, and +T3 were treated with 0.05% tetracycline. Gut-restoration lines +T11, +T22, and +T33 were treated with feces from the control lines in Generations 5 and 6.

### Antibiotic treatment

The antibiotic-treated lines (+T1, +T2, and +T3) were treated with tetracycline hydrochloride (lot number SLBQ2368V, Sigma-Aldrich, Germany) mixed in the malt food in a final concentration of 0.05% (Schneider *et al.*, 2013) for three generations (Figure 1). After the treatment with tetracycline, these lines were fed regular malt food for another three generations.

### Gut flora restoration

In Generation 5, each of the antibiotic-treated lines (+T1, +T2, and +T3) was divided to create gut-restoration lines, namely +T11, +T22, and +T33 (Figure 1). Each line to be restored was provided feces from its corresponding control line; for example, line +T11 was provided feces of line −T1. Mating cages with feces from the control lines that had been inhabited for one week by control lines and thus contained their feces were used for the gut-restoration lines as a restoration mating cage. Each gut-restoration line was provided feces for two weeks.

### Larval locomotion

In Generation 6, 20 larvae aged 5 days old were randomly collected from all lines for the locomotion experiment. Each larva was placed on a 55-mm petri dish filled with 2% agarose and placed on a light pad (A4 Light Box, M.Way, China). The order of the larvae was chosen randomly. The locomotion of each larva was recorded for three minutes using a Sony XR155 Full HD video camera (Sony, Japan). The total crawling distance (mm) and average speed (mm s^−1^) was analyzed with wrMTrck plugin version 1.04 (Nussbaum-Krammer *et al.*, 2015) in ImageJ version 1.53c (Schneider *et al.*, 2012) with slight modifications (Brooks *et al.*, 2016). The experiment was done over three days between 9.00 am and 12.00 pm and was identical to that in Detcharoen et al. (2020). The locomotion data of the larvae were analyzed using nested ANOVA function in R version 4.0.3 (R Core Team, 2020) with an alpha = 0.05 to identify significant differences within and between the lines or groups.

### Adult locomotion

Two methods were used for the adult locomotion experiments in Generation 6, the Rapid Iterative Negative Geotaxis (RING) (Gargano *et al.*, 2005) and the *Drosophila* Activity Monitor (DAM5M) (Trikinetics, USA). For each experiment, 20 2-week-old female flies of each line were randomly selected, anesthetized with CO_2_ for sexing three days before the experiment, and were put into separate vials with malt food. For both methods, flies were acclimatized at room temperature (19 °C) for one hour before the start of the experiments. The experiments took place between 9.00 am and 1.00 pm.

For the RING experiment, the flies were transferred individually into heptane-cleaned vials (100 x 24 x 1 mm, Scherf-Präzision Europa, Germany) and clamped in random order in the RING apparatus. The fly-containing vials were tapped quickly on the table, and the locomotion activities (walking and jumping) of flies were video recorded for three minutes with a video camera (Sony XR155 Full HD video camera, Sony, Japan). All fly lines were included in every run. The recorded videos were analyzed with ImageJ version 1.53c (Schneider *et al.*, 2012). The instances of locomotion activities were counted manually. The experiment was identical to that in Detcharoen et al. (2020).

For the second locomotion experiment, each fly was transferred into a glass vial and placed in the DAM5M in a random order. All fly lines were included in every run. The activities were measured every 20 seconds for an hour. The numbers of moves were detected automatically once a fly cross the infrared beam. The recorded data were then checked with DAMFileScan111X version 1.11 (Trikinetics, USA). The statistical analyzes were the same as for the larvae.

### Wing geometric morphometrics

To demonstrate any difference of tetracycline on wing morphology, 20 2-week-old female flies per line at Generation 6 were used. Wings were removed from each fly and stored in 96% ethanol. The upper and the lower side of left and right wings of each fly were photographed using a Leica Z6 APO macroscope with a Leica MC190 HD camera with 2.0x objective lens using to the Leica Application Suite version 4.0 (Leica Microsystems, Switzerland). The wing photos were converted to a tps file using tpsUtil32 version 1.79 (https://life.bio.sunysb.edu/morph/soft-utility.html). Thirteen landmarks were digitized manually using tpsDIG2w32 version 2.31 (https://life.bio.sunysb.edu/morph/soft-dataacq.html) on every photo. The wing photos with the landmarks were analyzed with MorphoJ version 1.07a (Klingenberg, 2011). The landmarks were aligned by the principal axis, and outliers were removed manually. Using Procrustes ANOVA, the potential imaging error between the lower and upper sides of the wing was accessed. Discriminant analysis between wings of all groups was performed. Canonical variate analyses (CVA) with 10,000 permutations was performed using regression residuals between centroid size and Procrustes coordinates. Principal component 1 and 2 of principal component analysis were exported to R to calculate Analysis of similarity (ANOSIM) between groups using an R package vegan version 2.5-6 (Oksanen *et al.*, 2019).

### Microbiome

Ten randomly chosen 14-day-old female flies were used per line. The antibiotic-treated lines were examined for a first time in Generation 2, and all the lines (control, antibiotic-treated, and gut-restoration) were checked in Generation 6. Each fly was killed in liquid nitrogen, surface sterilized using 2.5% bleach, and washed twice with sterile MilliQ-water (Chandler *et al.*, 2011), and the gut (crop, midgut, and hindgut) was removed under a stereo microscope (SMZ800, Nikon, Japan) on a sterile slide with sterile forceps. Five guts from the same line per replicate were transferred into a sterile 1.5 ml microcentrifuge tube, two replicates per line. The guts were homogenized manually with sterile plastic pestles. Mock microbial cells and DNA community standards (Zymo Research, USA) were used to check for DNA extraction efficiency and sequencing errors, respectively. One blank sample was used to check for bacterial contamination in the DNA extraction kit. DNA was extracted with QIAamp DNA Mini Kit (QIAGEN, Hilden, Germany). The DNA was resuspended in sterile water. Human DNA contamination was checked using Alu J primer (Cannas *et al.*, 2009) with quantitative PCR. The extracted DNA was amplified with bacterial 16S V3-V4 region of ribosomal DNA universal primer 341F and 805R (Herlemann *et al.*, 2011). The samples were sequenced with Illumina MiSeq 2 x 300 bp using a single lane at IGA Technology Services (Udine, Italy). The Qiime2 pipeline version 2020.6 (Bolyen *et al.*, 2019) was used for sequence analyses. DADA2 (Callahan *et al.*, 2016) implemented in Qiime2 was used to trim the sequences, merge forward and reverse reads, and remove chimeras. SILVA release 138 (Quast *et al.*, 2013) was used for a taxonomic assignment. Alpha and beta diversity were analyzed. ANOSIM on the phyla level was used to check the dissimilarity between groups and generations. The Kruskal-Wallis test was used to check whether independent samples came from a common group (control, antibiotic-treated, and gut-restoration). Non-metric Multi-dimensional Scaling (NMDS) at the phylum level was used to observe similarity of the different lines. Differential abundance and significant differences at the bacterial species level between the three groups was calculated using DESeq2 version 1.30.0 (Love *et al.*, 2014) in R.

## RESULTS

### Locomotion

The larvae of the gut-restoration line +T33 moved the furthest (mean 6.73 cm ± standard error 0.34 cm) (Figure 2A), and −T3 moved the fastest (0.038 cm s^−1^ ± 0.004 cm s^−1^) (Figure 2B). However, there were no significant differences among groups regarding mean speed and distance (nested ANOVA, speed, F_2,6_ = 0.12, p = 0.88; distance, F_2,6_ = 0.12, p = 0.88). We observed significant differences within the control group in crawling distance and speed (ANOVA, speed, F_2,57_ = 11.47, p < 0.001; distance, F_2,57_ = 12.13, p < 0.001), and significant difference in speed within the antibiotic-treated group (ANOVA, speed, F_2,57_ = 3.96, p = 0.02) (Figure 2B).

**Figure 2.**
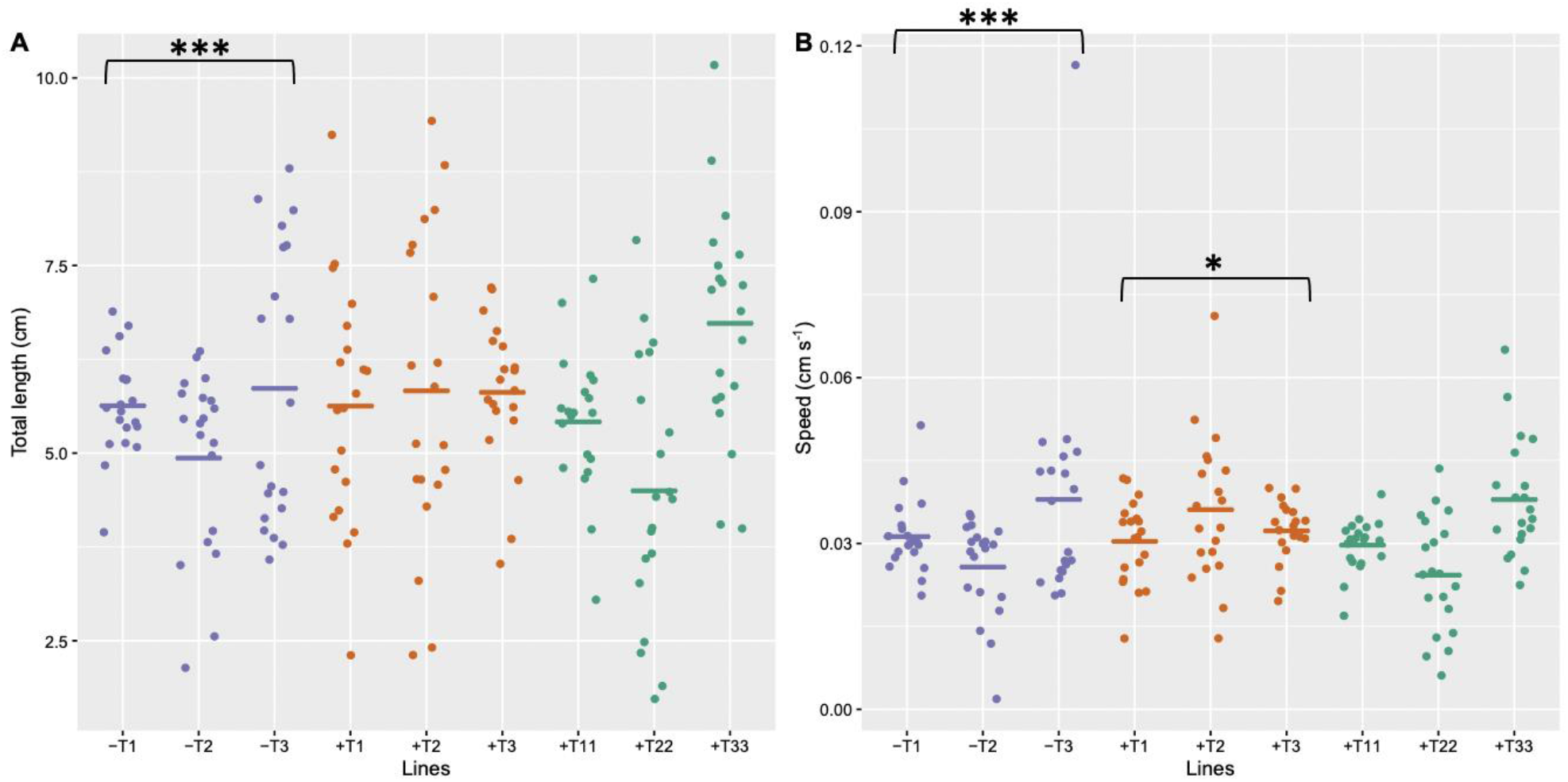
Each dot represents total length (A) and speed (B) of movements of larvae of *Drosophila nigrosparsa* in three minutes (N=20 for each line). Control (−T1, −T2, and −T3), antibiotic-treated (+T1, +T2, and +T3), and gut-restoration group (+T11, +T22, and +T33) are shown in purple, orange, and green, respectively. Plots show different y-scales. p < 0.001 is shown as ***. p < 0.05 is shown as *.

In the RING locomotion, the antibiotic-treated group had the highest walk (mean 3.57 walks ± standard error 0.42 walks) and jump (0.60 jumps ± 0.17 jumps) activities (Figure 3AB). However, there were no significant differences among the groups in terms of walk and jump activities (nested ANOVA, walk, F_2,6_ = 0.99, p = 0.42; jump, F_2,6_ = 0.21, p = 0.81).

**Figure 3.**
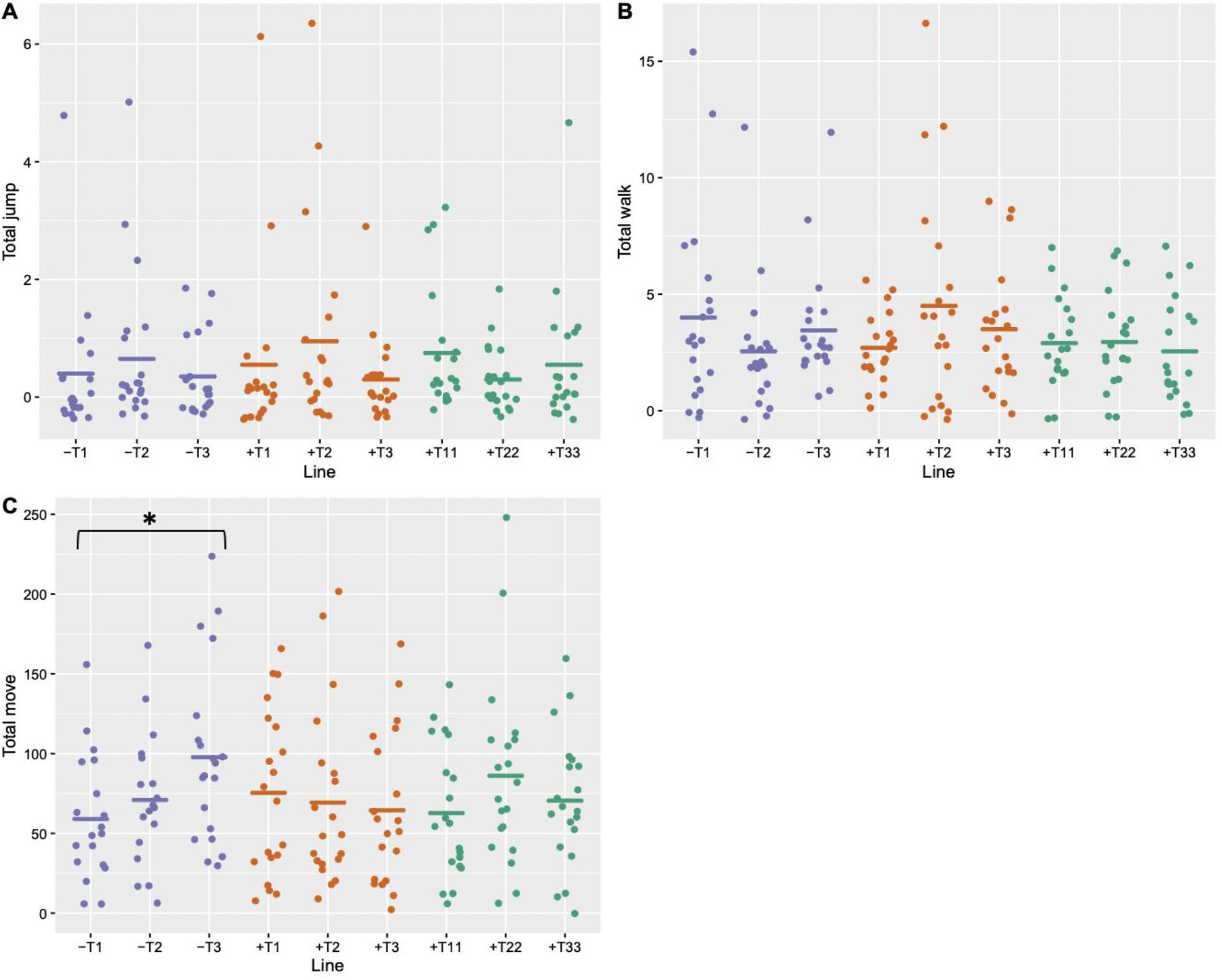
Jump (A) and walk (B) activities of adult female *Drosophila nigrosparsa* from the rapid iterative negative geotaxis experiment. Number of moves (C) of adult female *D. nigrosparsa* using the DAM5M method. Control (−T1, −T2, and −T3), antibiotic-treated (+T1, +T2, and +T3), and gut-restoration group (+T11, +T22, and +T33) are shown in purple, orange, and green, respectively (N = 20 for each line). Plots show different y-scales. P < 0.05 is indicated by ^*^.

In the DAM5M locomotion, the control group moved the most (mean 76 moves ± standard error 6 moves), especially line −T3 (97 moves ± 12 moves) (Figure 3C). There was no significant difference among groups (nested ANOVA, F_2,6_ = 0.17, p = 0.84), but there were significant differences within the control group (ANOVA, F_2,54_ = 3.58, p = 0.03).

### Geometric morphometrics

The mean squares of imaging error were very low for both centroid size and shape (46.6 and 5.7 times lower than individual by side interactions for centroid size and shape, respectively). There were significant differences among the fly lines in terms of shape and centroid size (Procrustes ANOVA; shape, F_44,154_ = 10.1, p < 0.001; size, F_2,7_ = 7.07, p < 0.001). After removing 6.3% of total variation within groups, calculated from regression, canonical variate analysis showed a slight grouping of the three groups (Figure 4). The Mahalanobis distance between control group and antibiotic-treated group was 1.72 (p < 0.001), between control group and gut-restoration group was 2.07 (p < 0.001), and between antibiotic-treated group and gut-restoration group was 1.62 (p < 0.001). ANOSIM static R value between antibiotic-treated and control groups was less than 0.01 (p = 0.45), between antibiotic-treated and gut-restoration groups was 0.02 (p = 0.05), and between control and gut-restoration groups was less than 0.01 (p = 0.77).

**Figure 4.**
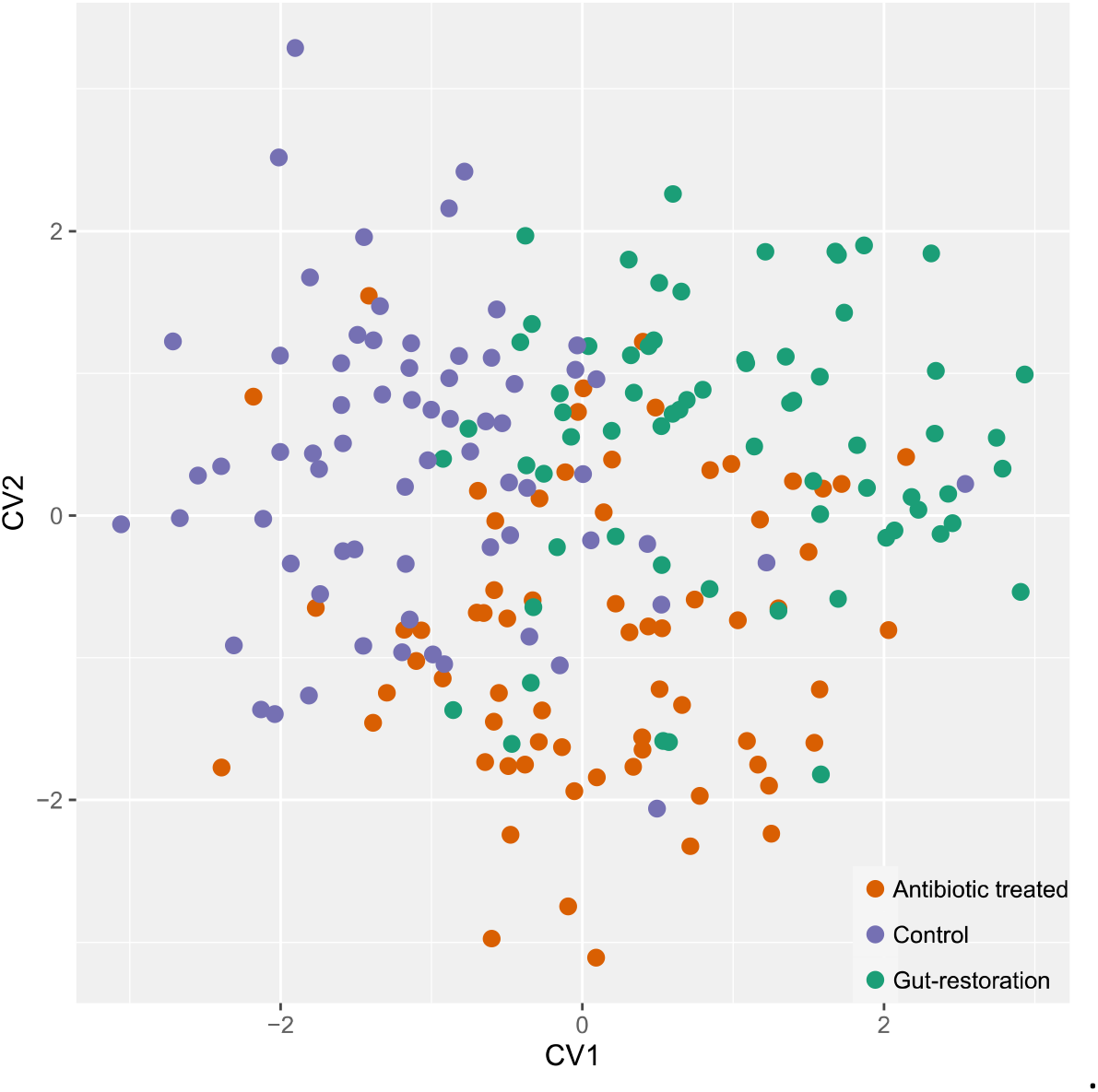
Canonical variate analysis with 10,000 permutations of wings of control (−T1, −T2, and −T3) (purple), antibiotic-treated (+T1, +T2, and +T3) (orange), gut-restoration (+T11, +T22, and +T33) (green) group. Each dot represents an individual *Drosophila nigrosparsa* female at Generation 6 (N=20).

### Microbiome

After trimming, the forward and the reverse sequences were 280 and 220 bases, respectively. A minimum merged read was 37,034 from a blank sample and a maximum of 390,048 reads from −T3 Generation 6 sample. The cell and DNA mock communities revealed a minor extraction and sequencing error (Figure S1). The mock cell extraction deviated in total 5% and 1% from the relative abundance from the mock cell and mock DNA community standards, respectively. The sequences found in the blank control were removed from all other samples.

Absolute abundances of sequences of antibiotic-treated samples at Generation 2 were lower than those of the other groups (Figure 5D). At the phylum level, all samples from Generation 6 were dominated by Proteobacteria and Firmicutes dominated the samples from the antibiotic-treated groups from Generation 2 (Figure 5B). At the genus level, *Acetobacter* was dominant in Generation 6. Line +T33 had the highest relative abundance of *Acetobacter* with 98.3%, followed by +T2 (97.5%) and +T1 (97.2%) (Figure 5B). The pairwise Kruskal-Wallis test revealed a significant difference between the control and gut-restoration groups (p = 0.05) but neither between the control and the antibiotic-treated group (p = 0.26) nor between the antibiotic-treated and the gut-restoration group (p = 0.26). NMDS at phylum level showed a separation between the samples from Generation 2 and the samples from Generation 6, supported by ANOSIM R value = 0.91 (p < 0.001) (Figure 6A). Comparing among groups at the phylum level, we found significant differences between the antibiotic-treated lines from Generations 2 and 6 (ANOSIM, R = 0.86, p = 0.004), but not between the gut-restoration and control lines (R = 0.04, p = 0.25), the gut-restoration and antibiotic-treated lines (R = 0.02, p = 0.29), and the antibiotic-treated and control lines (R = −0.05, p = 0.61). Differential abundance analyses at species level, *Acetobacter malorum* was significantly different in Generation 6 between the control group and the antibiotic-treated group and between the control group and the gut-restoration group together with another seven bacterium species (Table 1). No significant differences in taxa abundance between antibiotic-treated and gut-restoration group were found.

**Table 1.**
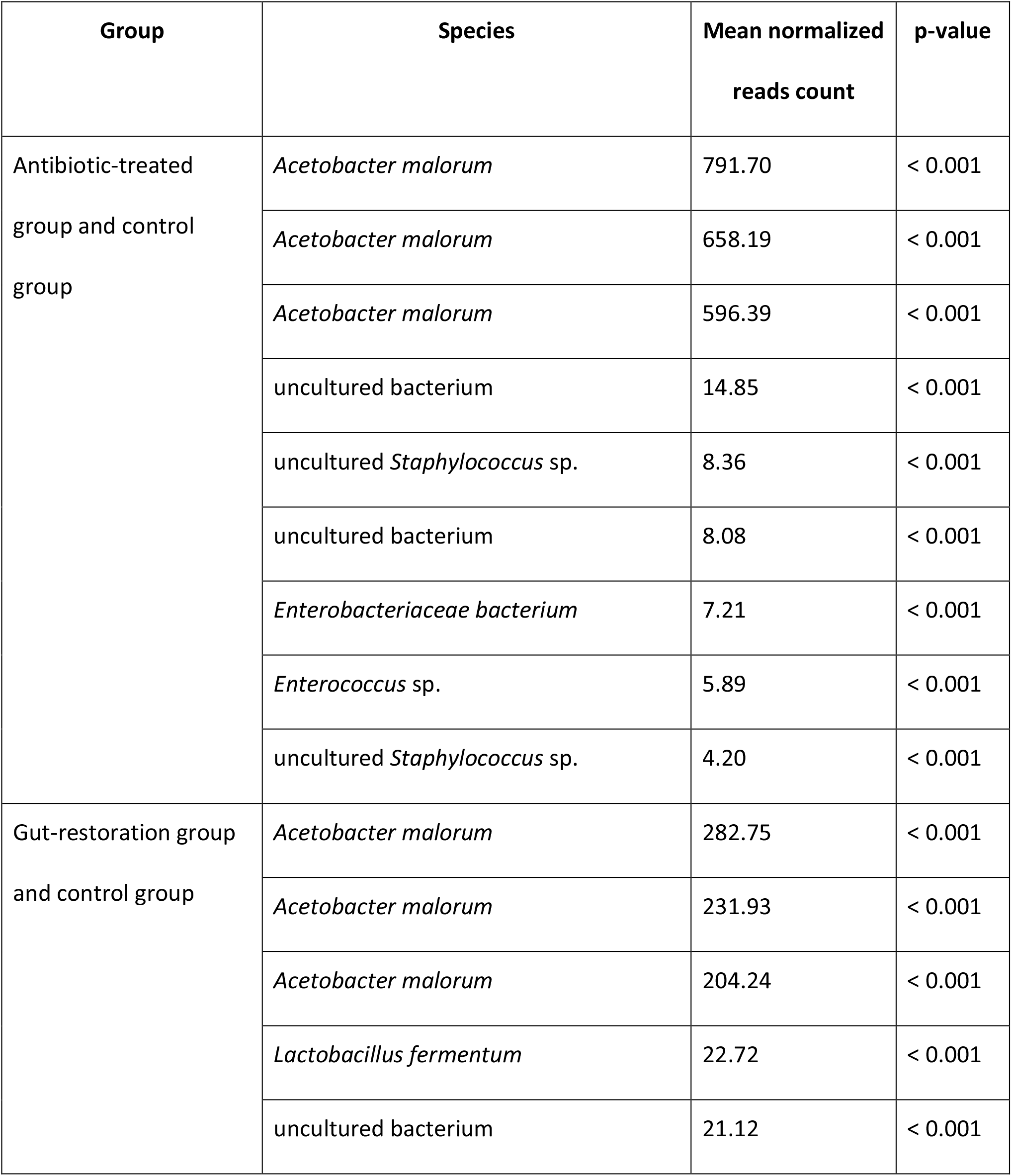
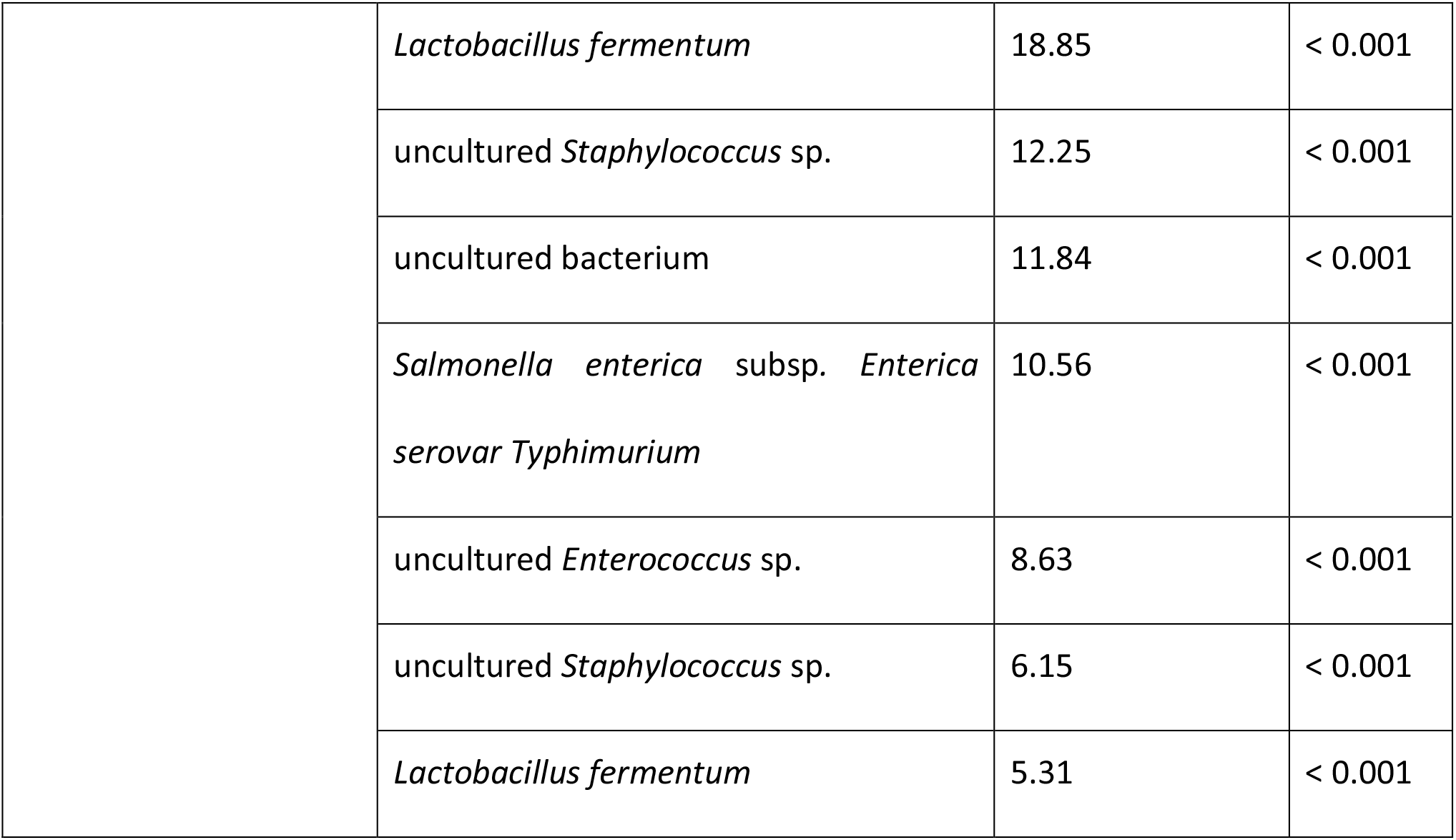
Differential abundance of bacterial species between the control group and the antibiotic-treated group and between the control group and the gut-restoration group. No species was significantly different between the gut-restoration group and the antibiotic-treated group.

**Figure 5.**
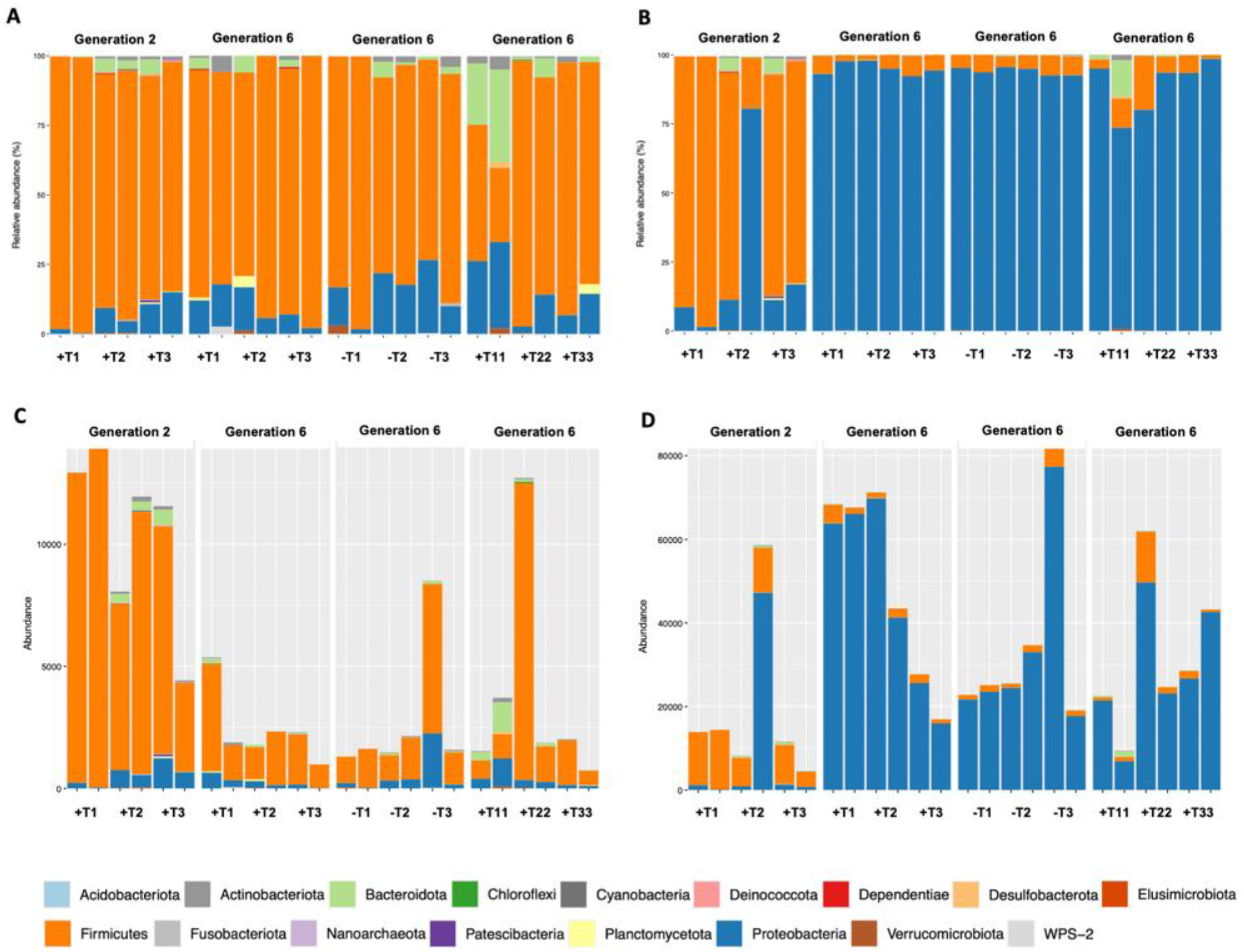
Relative (A, B) and absolute abundance (C, D) of bacterial phyla calculated without (A, C) and with *Acetobacter* species (B, D). Two replicas per line are shown. Antibiotic-treated group Generation 2 and 6 (+T1, +T2, and +T3), control group Generation 6 (−T1, −T2, and −T3), and gut-restoration group Generation 6 (+T11, +T22, and +T33) are shown. Plots have different y-scales.

**Figure 6.**
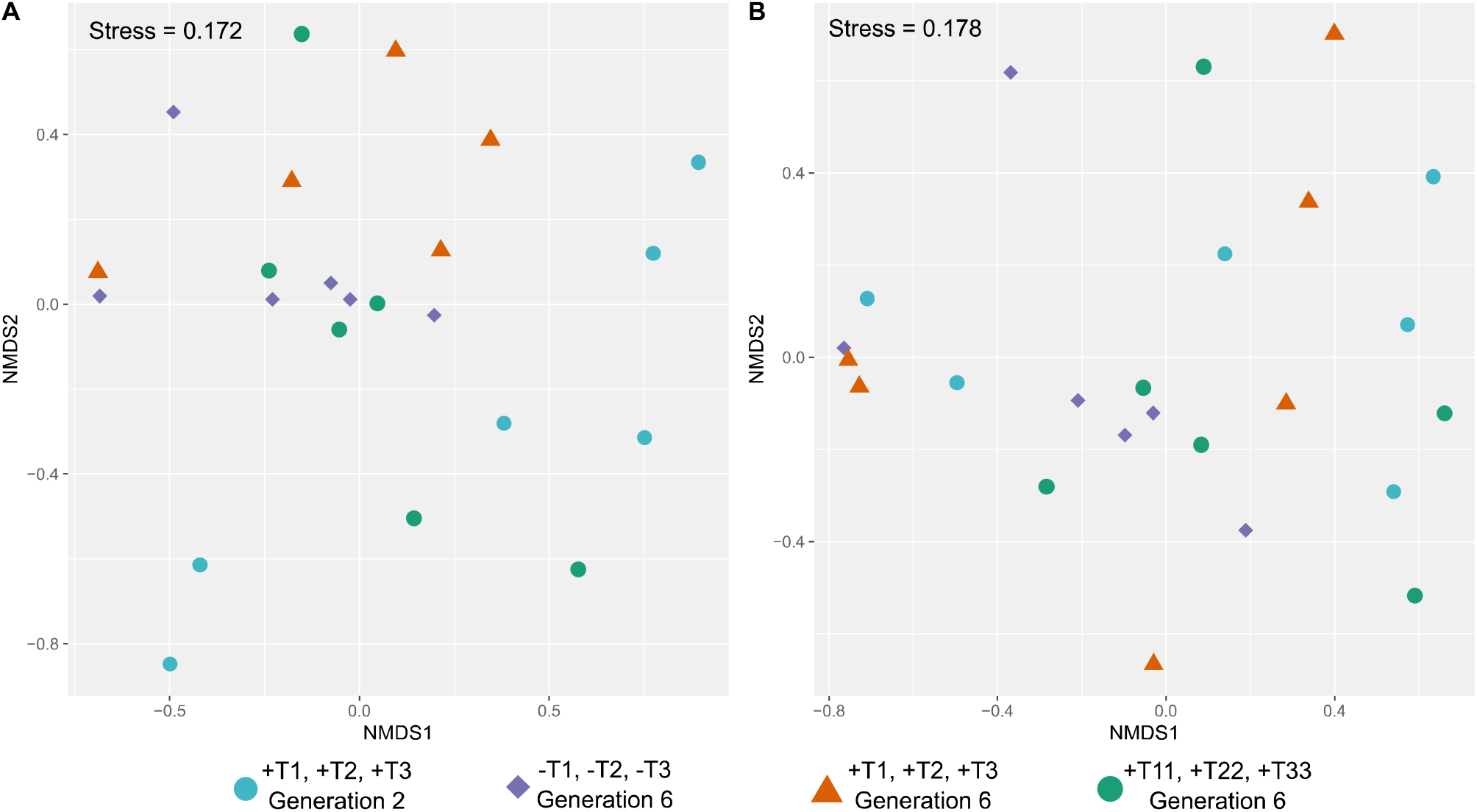
Non-metric multidimensional scaling of samples at phylum level with (A) and without *Acetobacter* spp. (B). Antibiotic-treated group Generation 2 (turquoise) and Generation 6 (orange), control group Generation 6 (purple) and gut-restoration group Generation 6 (green). Plots show different x/y-scales.

To analyze the other bacterial phyla in detail, sequences of *Acetobacter* were removed. The phylum Firmicutes dominated with 99.3% of line +T1 at Generation 2, followed by −T1 (98.3%) and +T3 (98.0%), all from Generation 6 (Figure 5A). No significance among groups emerged from the pairwise Kruskal-Wallis test (control and antibiotic-treated group: p = 0.26; control and gut-restoration group: p = 0.87; gut-restoration and antibiotic-treated group: p = 0.77). The exclusion of *Acetobacter* resulted in a more compact arrangement of all groups (Figure 6B). No significant difference found between Generation 2 and 6 (ANOSIM, R = −0.12, p = 0.83), within antibiotic-treated lines from Generation 2 and 6 (R = −0.13, p = 0.92), between control and gut-restoration line (R = −0.03, p = 0.59), and antibiotic-treated and control line (R = −0.04, p = 0.53).

## DISCUSSION

The gut-restoration protocol with two 7-day restoration cycles from the respective control line (−T1, −T2, and −T3) worked well in practice. Subjectively, no abnormality (e.g., increased mortality or decreased fecundity) in the general condition of the acceptor groups (+T11, +T22, and +T33) was identified. However, no data were collected in this regard.

Larval locomotion did not differ among the three groups, control, antibiotic-treated, and gut-restoration. Instead, we found variation among lines within groups, that is, significant differences in mean speed and total distance within the control group and a significant difference in mean speed within the antibiotic-treated group (Figure 2). In addition, we did not find significant differences among the three groups in neither of the two adult locomotion experiments RING and DAM5. These locomotion results indicate that two generations recovering time after tetracycline treatment are sufficient to eliminate side effects regarding larval locomotion. As we did not find any significant difference between control and antibiotic-treated group, we can confirm that our previous results on locomotion that *Wolbachia*-infected flies had higher locomotion activities than antibiotic-treated flies (Detcharoen *et al.*, 2020) were due to effects of *Wolbachia*. However, changes in locomotion like walking speed and pattern can be induced by gut microbiome (Schretter *et al.*, 2018).

We found a slight grouping of the fly wings into the three corresponding treatment groups (Figure 4). One of the reasons for the slight grouping of our samples could be genetic drift. The effects of drift can be observed after a few generations after separation of flies, like in previous studies in *D. nigrosparsa* (Detcharoen *et al.*, 2020) and *Drosophila subobscura* (Santos *et al.*, 2013). However, genetic drift appears unlikely here, as not only the groups but also the lines have been separated for six generations (Figure 1). Although genetic variation is highly reduced in *Drosophila* isofemale lines, morphological differences can persist (Bubliy *et al.*, 2001; Carreira *et al.*, 2006). One reason for the slight morphological changes in the wings could be differences in the microbiome, like was shown for gut morphology of *D. melanogaster* (Broderick *et al.*, 2014).

Firmicutes was the most dominant bacterial phylum of all samples of the antibiotic-treated group in Generation 2, whereas this phylum became the second most dominant phylum in Generation 6, after Proteobacteria (Figure 5B D). A decrease in Proteobacteria and an increase in Firmicutes during antibiotic treatment was observed in the earthworm *Metaphire guillelmi* treated with tetracycline (Chao *et al.*, 2020). Proteobacteria of the soil microbiome decreased rapidly when treated with tetracycline (Lin *et al.*, 2016). However, tetracycline treatment does not always lead to increase in Firmicutes, for example, in the small brown planthopper *Laodelphax striatellus* where the abundance of firmicutes bacterial decreased (Zhang *et al.*, 2020). Recently, a study found that phyla Proteobacteria (genus *Acetobacter*) and Firmicutes were associated with short-term memory, learning, and cognition in the fly *D. melanogaster* (DeNieu *et al.*, 2019).

*Acetobacter* spp. were the most abundant in Generation 6. The significant in differential abundance of bacterial taxa we found between the control group and the antibiotic-treated group in Generation 6 (Table 1) means that two generations after the last antibiotic treatment were not enough for gut microbiome to recover. One should consider alternative methods for gut microbiome restoration after tetracycline treatment.

We cannot make a clear statement regarding the potential influence of the microbiome on wing morphology. At first glance, the lack of significant difference in bacterial species between the gut-restoration group and the antibiotic-treated group speaks against it, as morphology of these groups does differ slightly. Along the same lines, the lack of significance in the ANOSIM calculations and the lack of groupings in the NMDS (Figure 6) can be interpreted as not in support of a direct influence of the microbiome. However, even if there is no linear effect discernible between the microbiome and the wing morphology, it can still be there.

Robust inferences from the combined evidence are that tetracycline had no significant influence on locomotion after two generations recovery time but did have such influence on the gut microbiome. These differences in gut microbiome among groups may have triggered a slight difference among groups in wing morphology, which would then be an indirect effect of the antibiotic. Together with the absence of an effect on locomotion, this suggests that checking for both direct and indirect effects of tetracycline after a particular recovery time before using tetracycline curing is important.

## ACKNOWLEDGMENTS

We thank Anja Ekblad for help with fly maintenance, Wolfgang Arthofer for his participation in initial discussions, and the University of Innsbruck for providing a doctoral grant to MD.

## CONTRIBUTION OF AUTHORS

B.C.S-S. and F.M.S. conceived this study with contributions from S.O.W. and M.D.; S.O.W., and M.D. collected and analyzed the data; S.O.W. drafted the paper with input from M.D., B.C.S-S., and F.M.S.

## DATA AVAILABILITY STATEMENT

Sequences are available on GenBank (BioProject accession number PRJNA694538). All other data will be made available in a public repository as soon as the manuscript has been accepted by the journal.

---

**Figure S1.**
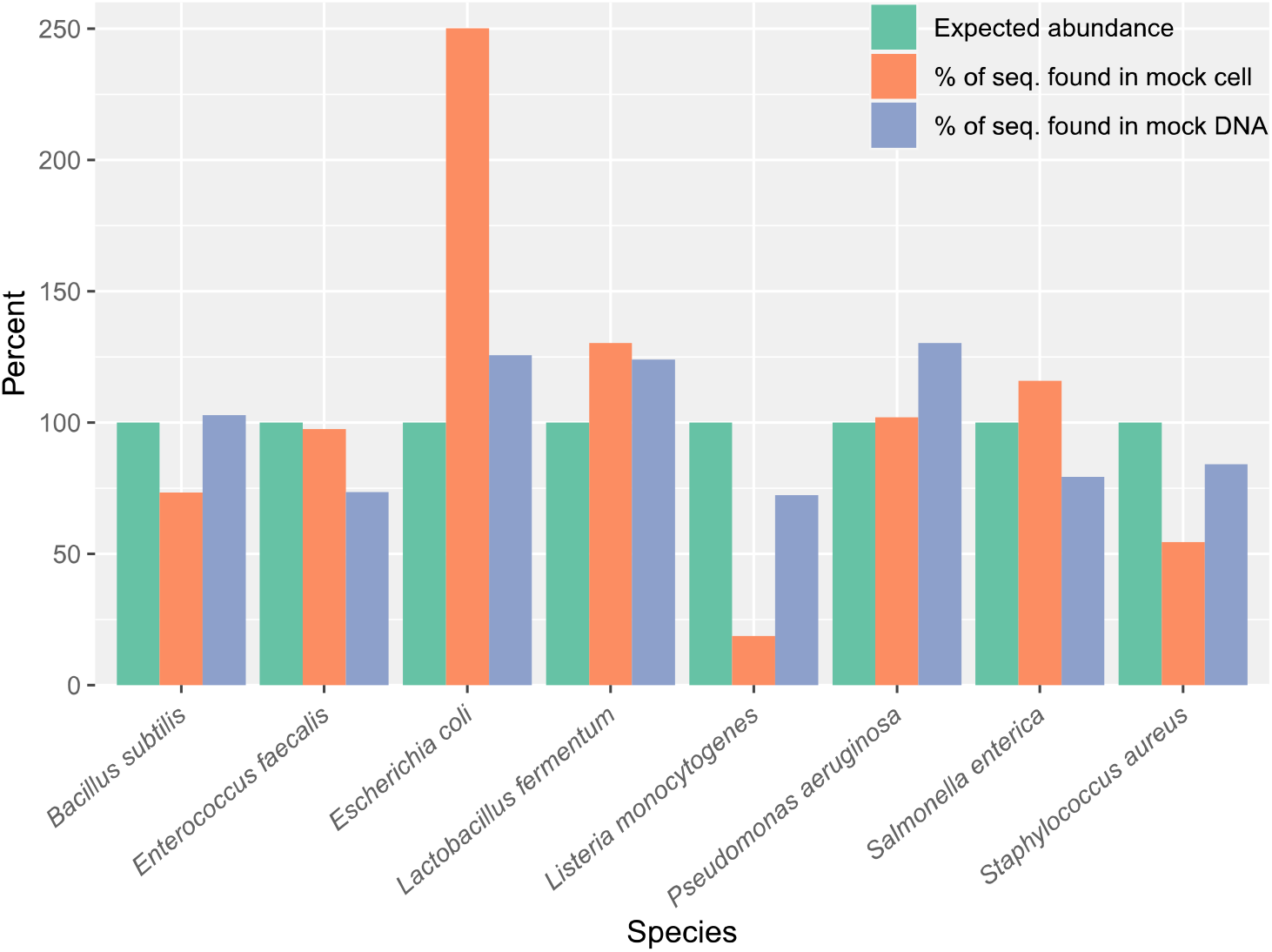
Positive control with percentage of sequences of mock cell (orange) and DNA (gray) communities (ZymoBIOMICS™, USA) from 16S sequencing compared with the expected results.

