## Supplementary figures and images for "No influence of antibiotic on locomotion in *Drosophila nigrosparsa* after recovery, but influence on microbiome, possibly mediating wing-morphology change"

### Supplementary figure S1

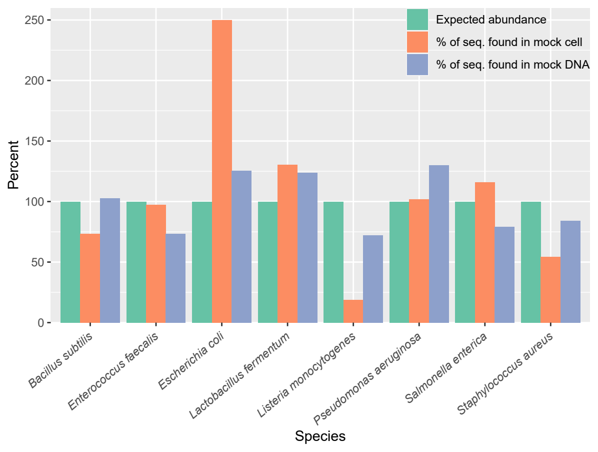
